# PconsC4: fast, free, easy, and accurate contact predictions

**DOI:** 10.1101/383133

**Authors:** Mirco Michel, David Menéndez Hurtado, Arne Elofsson

## Abstract

**Motivation:** Residue contact prediction was revolutionized recently by the introduction of direct coupling analysis (DCA). Further improvements, in particular for small families, have been obtained by the combination of DCA and deep learning methods. However, existing deep learning contact prediction methods often rely on a number of external programs and are therefore computationally expensive.

**Results:** Here, we introduce a novel contact predictor, PconsC4, which performs on par with state of the art methods. PconsC4 is heavily optimized, does not use any external programs and therefore is significantly faster and easier to use than other methods.

**Availability:** PconsC4 is freely available under the GPL license from https://github.com/ElofssonLab/PconsC4. Installation is easy using the pip command and works on any system with Python 3.5 or later and a modern GCC compiler.

## Introduction

To predict the structure of a protein from no other information than its sequence has been a major challenge in bioinformatics for decades. With the introduction of direct coupling analysis (DCA) to improve contact predictions (Weigt *et al.*, 2009) significant progress in protein structure prediction was reported in 2011 (Morcos *et al.*, 2011; Marks *et al.*, 2011). These methods have then been used to predict the structure of hundreds of protein families with high accuracy to an unprecedented accuracy (Ovchinnikov *et al.*, 2017). One disadvantage of DCA methods is that they require very large multiple sequence alignments to provide accurate contact predictions. This problem has been overcome by refining the initial DCA prediction with deep learning methods (Skwark *et al.*, 2014; Wang *et al.*, 2017).

Although the structure can accurately be predicted for many protein families, there still exist many families were the predictions are not sufficiently accurate. Although predictions are better for larger families, many other factors also seem to be important (Michel *et al.*, 2017a). The exact reason for what is needed to improve the predictions is not well understood. But it is not unlikely that the underlying multiple sequence alignments are not optimal. It might therefore be possible to improve the predictions if alternative multiple sequence alignments are examined or metagenomics data is included. Given the computational costs of earlier deep learning contact predictions methods it has been difficult to exhaustively examine alternative alignments for each protein family.

Here, we present a novel deep learning approach, PconsC4, that performs as well, or even better, than earlier methods. More importantly it is freely available, significantly faster and easier to install than alternative methods. This provides all users with an easy way to explore alternative multiple sequence alignments or other parameters for contact predictions.

## Implementation

Below follows a short description of PconsC4, for details see the supporting information. PconsC4 is trained on a set of 2791 proteins culled from PDB and benchmarked on two datasets without any homology to the training set. One validation set is identical to the benchmark set in PconsC3 (Michel *et al.*, 2017b). Additionally, we benchmarked PconsC4 on 44 proteins from CASP12, see Tables S2-S5. Multiple sequence alignments are created using three iterations of HHblits (Remmert *et al.*, 2011) and an e-value threshold of 1.0. Other alignment methods and cut-offs as well as combinations of different alignments were tested but did not provide significant improvements.

From each position in the multiple sequence alignments 72 features are calculated and fed into the PconsC4 network. These include: 68 one-dimensional sequential features and four pairwise features; the GaussDCA score (Baldassi *et al.*, 2014), APC-corrected mutual information, normalized APC-corrected mutual information, and cross-entropy.

At the core of PconsC4 is the U-net architecture (Ronneberger *et al.*, 2015), designed for image segmentation. It is composed of a series of convolutional layers, down- and up-sampling, with shortcut connections to help convergence.

To include secondary structure information but remain independent of external predictors, we took the pre-trained network from ProQ4 (Menendez *Hurtado et al.*, 2018), that takes all the onedimensional inputs and predicts secondary structure, dihedral angles, and surface accessibility for each residue. PconsC4 takes the output of the second to last layer, transforms them into twodimensional features via an outer product, and concatenates them to the rest of the inputs.

Finally, the network produces four outputs: the probability of a contact for the thresholds of 6, 8, or 10 A, and the distance measured as S-score.

To reduce the number of dependencies and overall run time, we re-implemented GaussDCA as a Python package using Pythran (Guelton *et al.*, 2015) resulting in a speedup of a factor of three. Optimization details are in section 2 of the supplementary information.

**Table 1.** Performance in PPV for the *L* top-ranked contact with a sequence separation of > 5 residues. Results for all, small families with Meff (Baldassi *et al,*, 2014) < and for short-, (5,12), medium- (12, 23) and long: (23,∞) ranges. Average runtime and external dependencies.

|  | All | Small | Short | Med | Long | Time [s] | Deps. |
| --- | --- | --- | --- | --- | --- | --- | --- |
| PlmDCA | 0.36 | 0.12 | 0.14 | 0.15 | 0.26 | 84 | - |
| GaussDCA | 0.34 | 0.12 | 0.14 | 0.15 | 0.25 | 27 | - |
| PSICOV | 0.34 | 0.11 | 0.13 | 0.14 | 0.22 | 114 | - |
| PhyCMAP | 0.32 | 0.29 | 0.24 | 0.19 | 0.18 | 718 | 1,3 |
| PconsC3 | 0.57 | 0.36 | 0.25 | 0.25 | 0.37 | 2931 | 1,2,3,4,5,6 |
| Metapsicov | 0.59 | 0.39 | 0.27 | 0.26 | 0.36 | 1158 | 1,3,7,8,9 |
| PconsC4 | 0.65 | 0.42 | 0.29 | 0.28 | 0.45 | 12 (56*) | - |

1: PSIPRED, 2: NetSurfP, 3: PSI-BLAST, 4: PhyCMAP, 5: PlmDCA, 6: GaussDCA, 7: PSICOV, 8: Freecontact, 9: CCMpred.
\* Timing without pre-loading the network.

## Results and Discussion

Table 1 shows a performance comparison of different methods running on the same alignments. PconsC4 performs 14% better than PconsC3 on the benchmark dataset and 10% better than Metapsicov. These improvements are consistent across short, medium, and long range contacts and for all thresholds (Figures S1-2). Additional evaluations are also presented in the supplementary information. As shown in Figures S3-4, the predicted scores are well calibrated, i.e. the reported scores reflect the real probability for a contact to exist. This enables easy comparison of alternative alignments.

All machine learning methods perform significantly better than the DCA methods for smaller families. PconsC4 approaches maximum performance at around 10^2^ effective sequences compared with 10^5^ for the DCA methods (Figure S5).

The overall performance is comparable to meta-meta predictors such as DNCON2 (Adhikari *et al.*, 2018) or other deep learning methods (Wang *et al.*, 2017). However, a direct comparison is difficult as these programs are either not available for download, to slow to run for large scale experiments, or cannot be used with a specific multiple sequence alignment.

The main advantage of PconsC4 is that it is fast and easy to use. Starting from a single input alignment PconsC4 predicts the contacts in less than one minute (or 12 s if the network is preloaded).

This is more than 25 times faster than PconsC3 or Metapsicov, while still consistently outperforming them. Since PconsC4 is not tied to any specific alignment method, it can directly take advantage of any improvement on this front, such as the use of alternative alignment strategies (Buchan and Jones, 2017) or metagenomics data (Ovchinnikov *et al.*, 2017), and thus be easily integrated into pipelines.

## Acknowledgements

Our thanks to Serge Guelton for his swift fixes to Pythran and to Nikos Tsardakas Renhuldt for discussion. This work was supported by grants from the Swedish Research Council (VR-NT 2016-03798 to AE). This research was conducted using the resources of High Performance Computing Center North (HPC2N).

## 1 Results

Here we report more detailed comparisons on the test sets. First, in Table S1 we compare short, medium, and long range contacts, showing that PconsC4 is better than previous methods at all ranges. Figures S1 and S2 show the receiver operating characteristic curves.

We also include an expanded version of the Table 1, Table S1 including CASP12 and other methods. For benchmarking GaussDCA we used the Julia implementation and CCMpred for PlmDCA. The times were measured on a machine with an Intel i7-4770 CPU and a NVIDIA 1070Ti GPU for CCMpred.

PconsC4 seems to be better calibrated that earlier methods, as shown in Figures S3 and S4. This means that a predicted score of 0.2 indicates that there is a 20% chance for a contact to exist. A perfectly calibrated predictor would lie on the diagonal (dotted line).

**Table S1.**
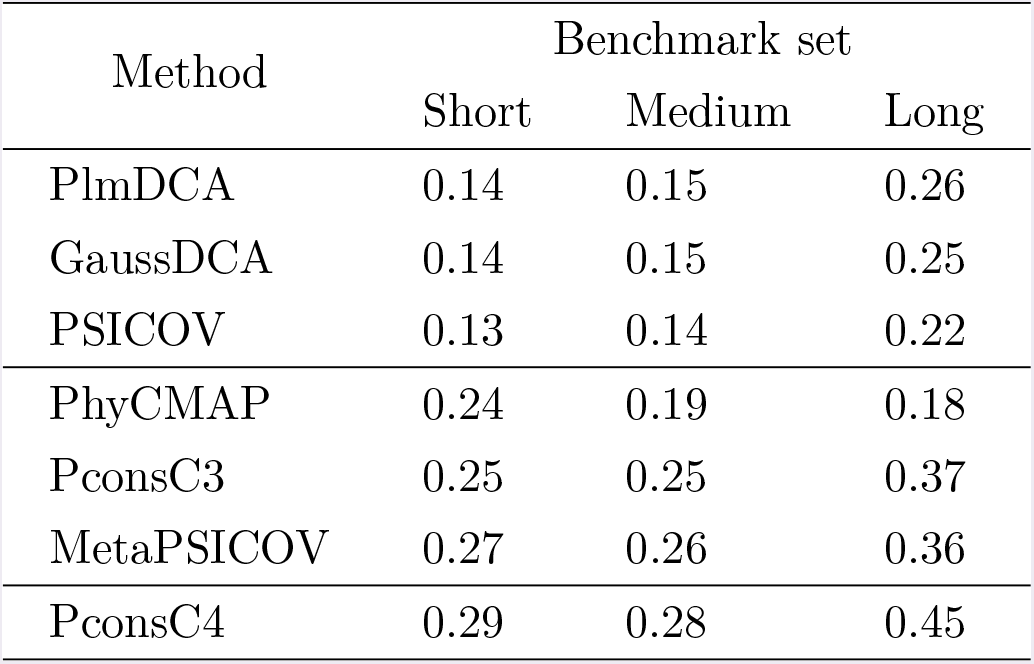
Precision for the top *L* contacts, where *L* is the sequence length, at different distance thresholds. Short: (5,12], medium: (12, 23], long: (23, ∞).

| Method | Benchmark set |  |  |
| --- | --- | --- | --- |
|  | Short | Medium | Long |
| PlmDCA | 0.14 | 0.15 | 0.26 |
| GaussDCA | 0.14 | 0.15 | 0.25 |
| PSICOV | 0.13 | 0.14 | 0.22 |
| PhyCMAP | 0.24 | 0.19 | 0.18 |
| PconsC3 | 0.25 | 0.25 | 0.37 |
| MetaPSICOV | 0.27 | 0.26 | 0.36 |
| PconsC4 | 0.29 | 0.28 | 0.45 |

**Table S2.**
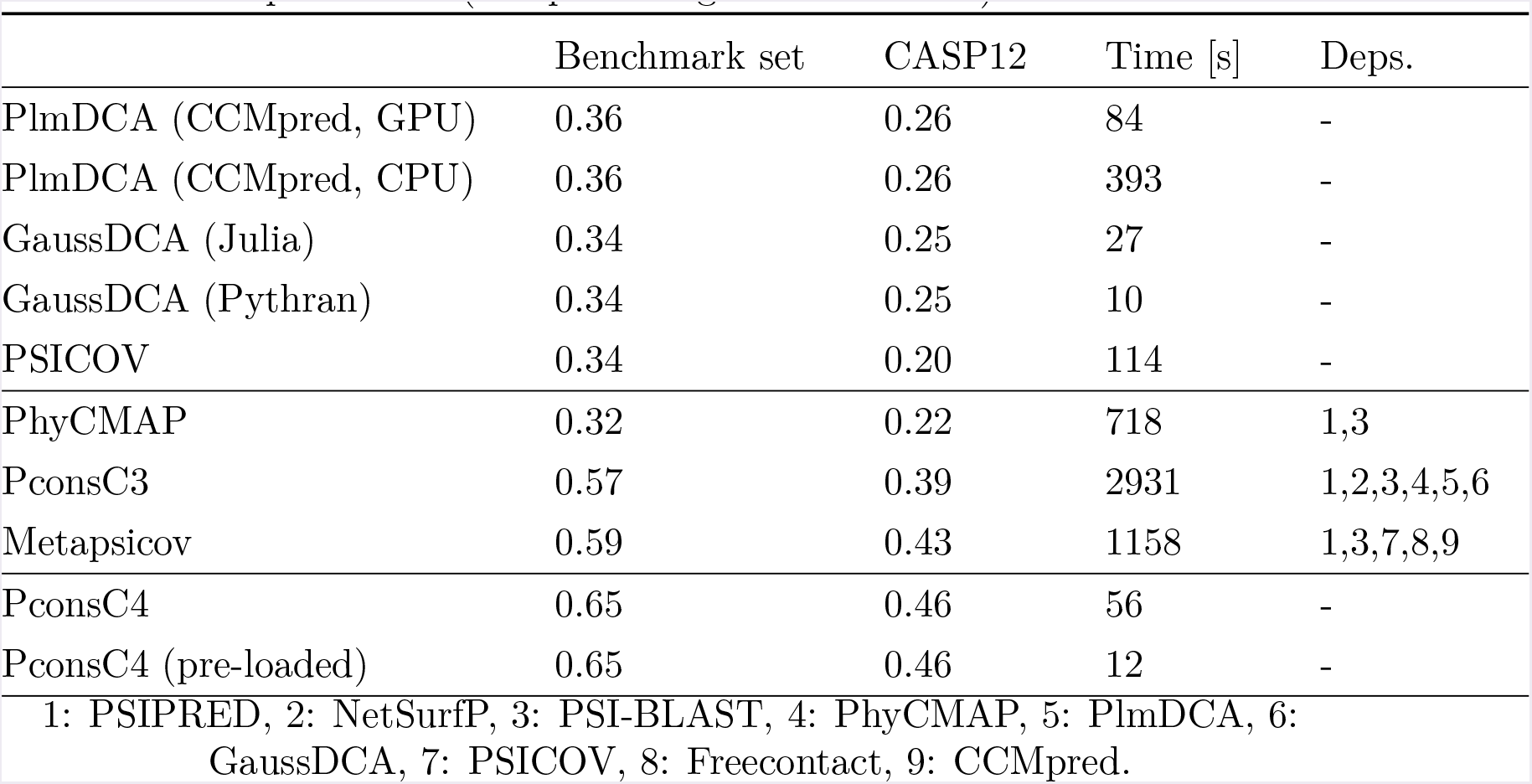
Performance in PPV for the *L* top-ranked contact with a sequence separation of > 5 residues, average runtime in seconds on the benchmark set, and external dependencies (except for alignment methods).

|  | Benchmark set | CASP12 | Time [s] | Deps. |
| --- | --- | --- | --- | --- |
| PlmDCA (CCMpred, GPU) | 0.36 | 0.26 | 84 | - |
| PlmDCA (CCMpred, CPU) | 0.36 | 0.26 | 393 | - |
| GaussDCA (Julia) | 0.34 | 0.25 | 27 | - |
| GaussDCA (Pythran) | 0.34 | 0.25 | 10 | - |
| PSICOV | 0.34 | 0.20 | 114 | - |
| PhyCMAP | 0.32 | 0.22 | 718 | 1,3 |
| PconsC3 | 0.57 | 0.39 | 2931 | 1,2,3,4,5,6 |
| Metapsicov | 0.59 | 0.43 | 1158 | 1,3,7,8,9 |
| PconsC4 | 0.65 | 0.46 | 56 | - |
| PconsC4 (pre-loaded) | 0.65 | 0.46 | 12 | - |

1: PSIPRED, 2: NetSurfP, 3: PSI-BLAST, 4: PhyCMAP, 5: PlmDCA, 6: GaussDCA, 7: PSICOV, 8: Freecontact, 9: CCMpred.

### 1.1 ROC and calibration curves

**Figure S1.**
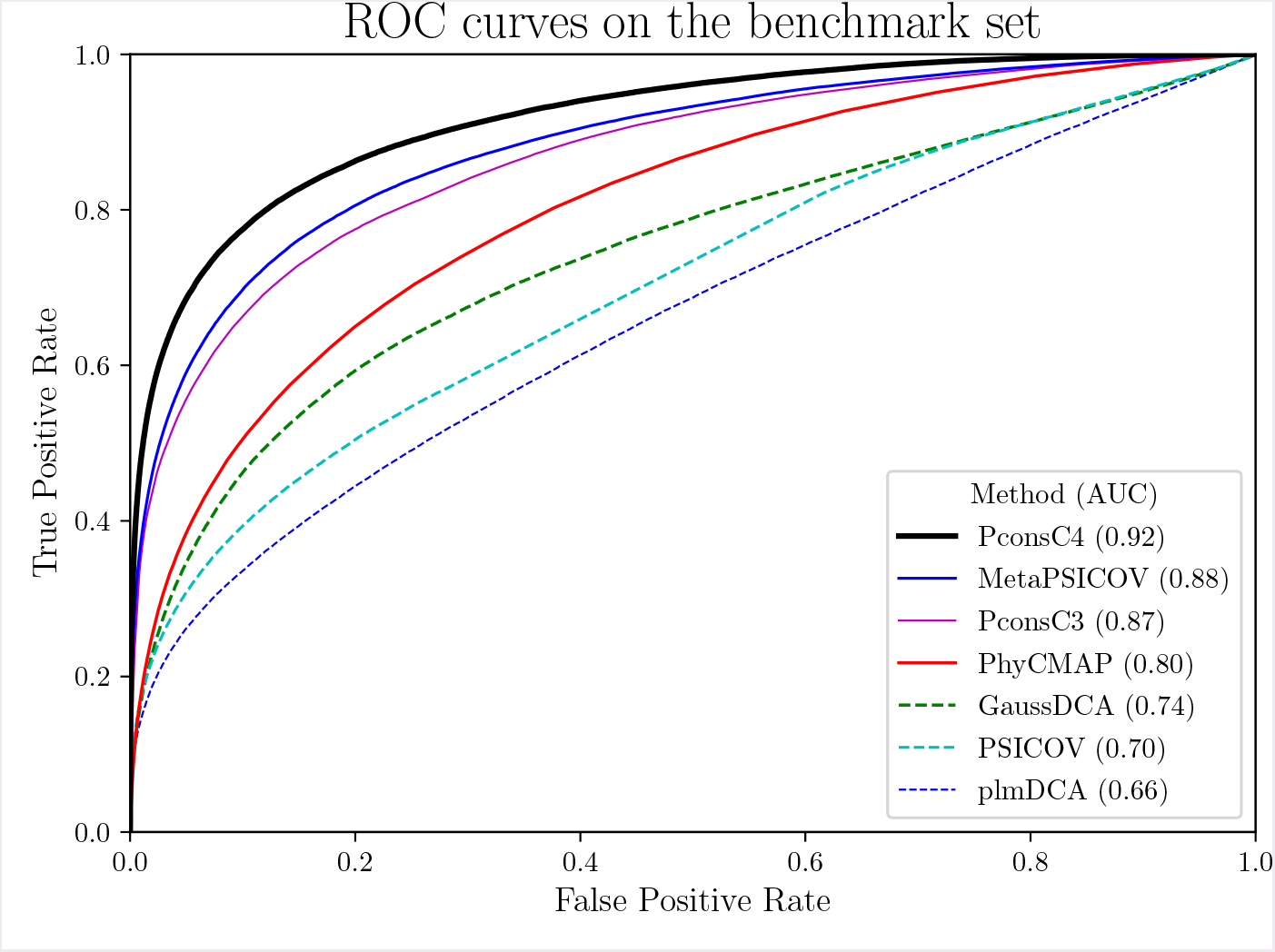
ROC curves on the Benchmark set

**Figure S2.**
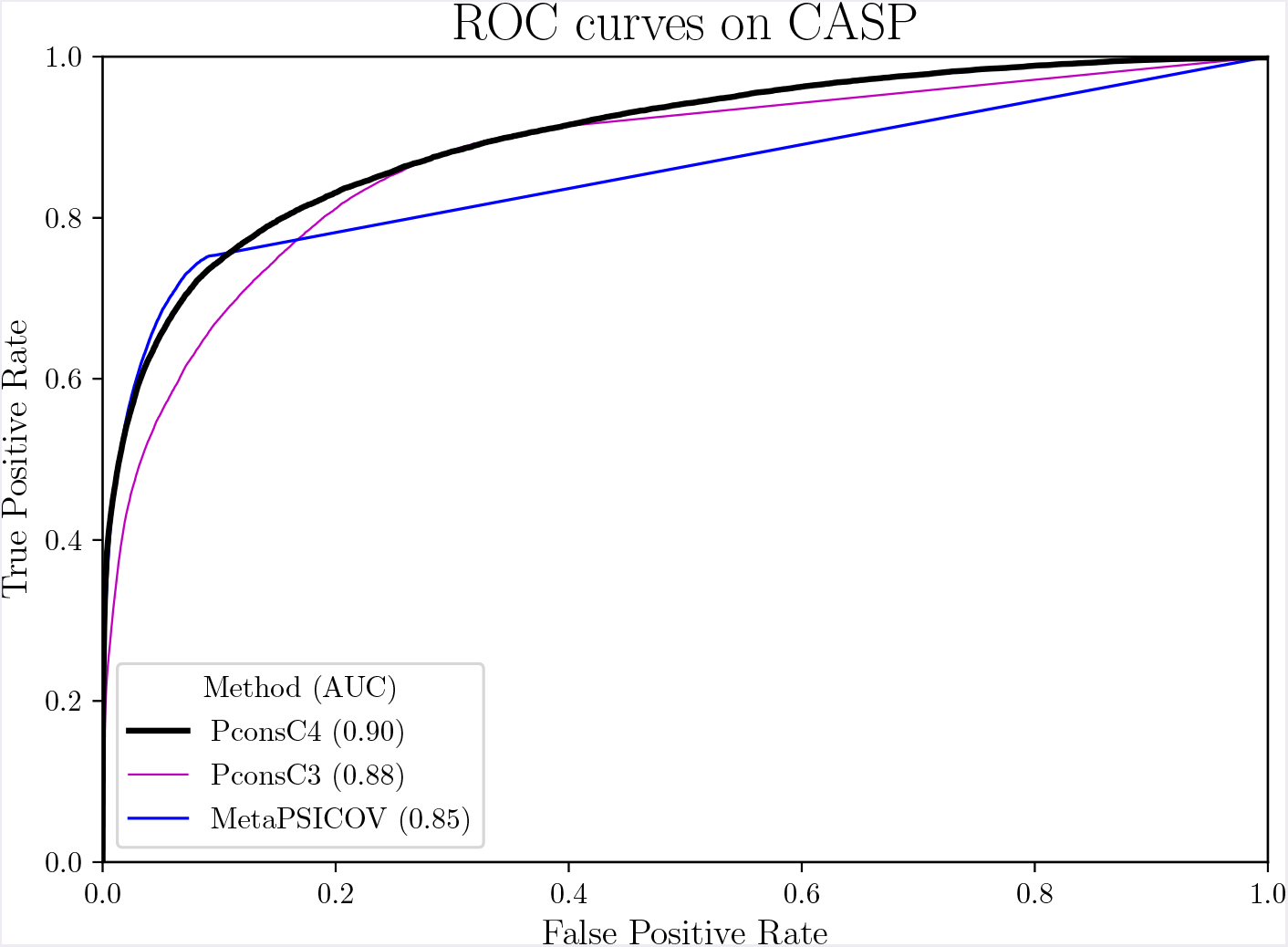
ROC curves on CASP12

**Figure S3.**
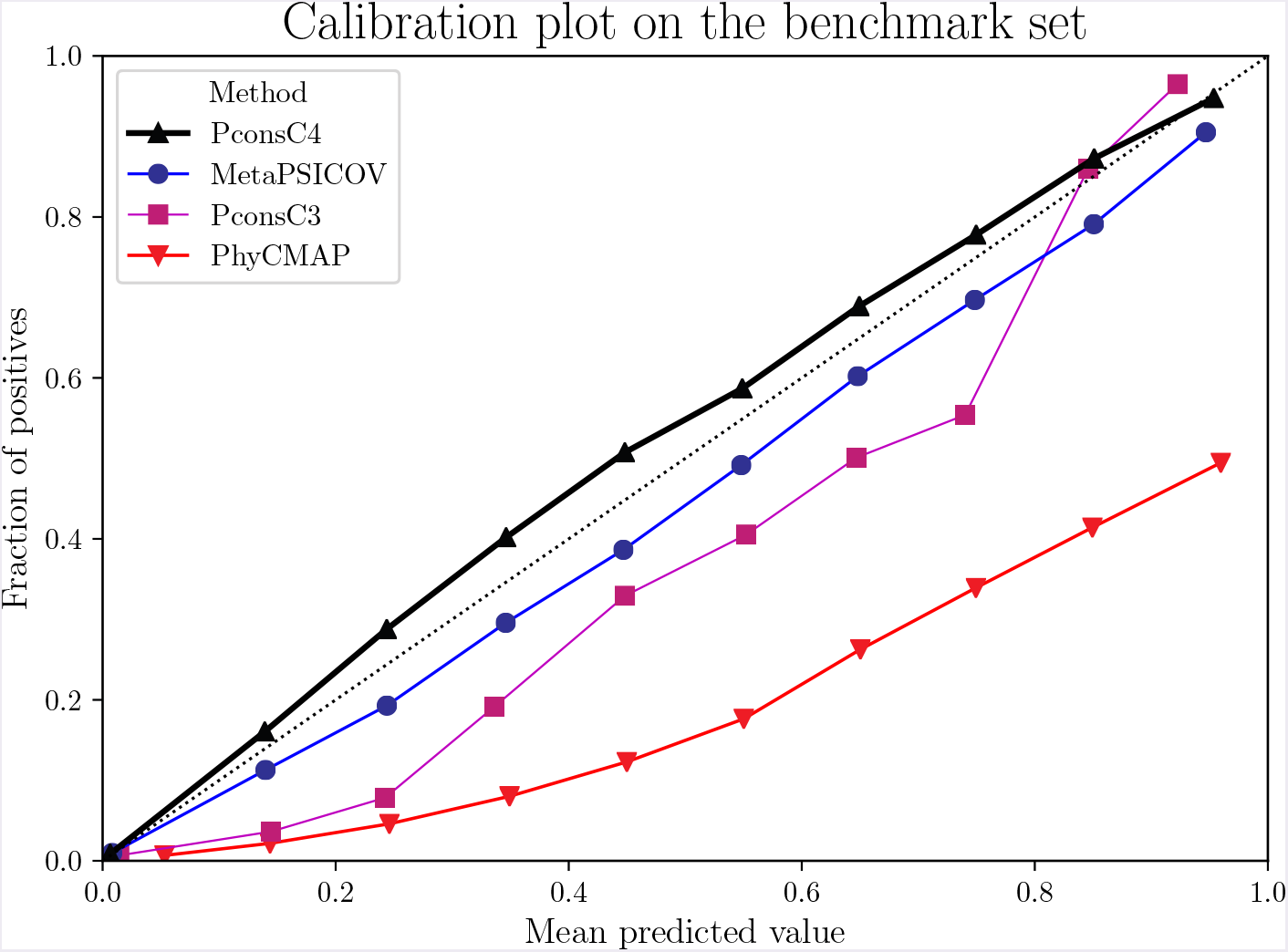
Calibration curves on the Benchmark set

**Figure S4.**
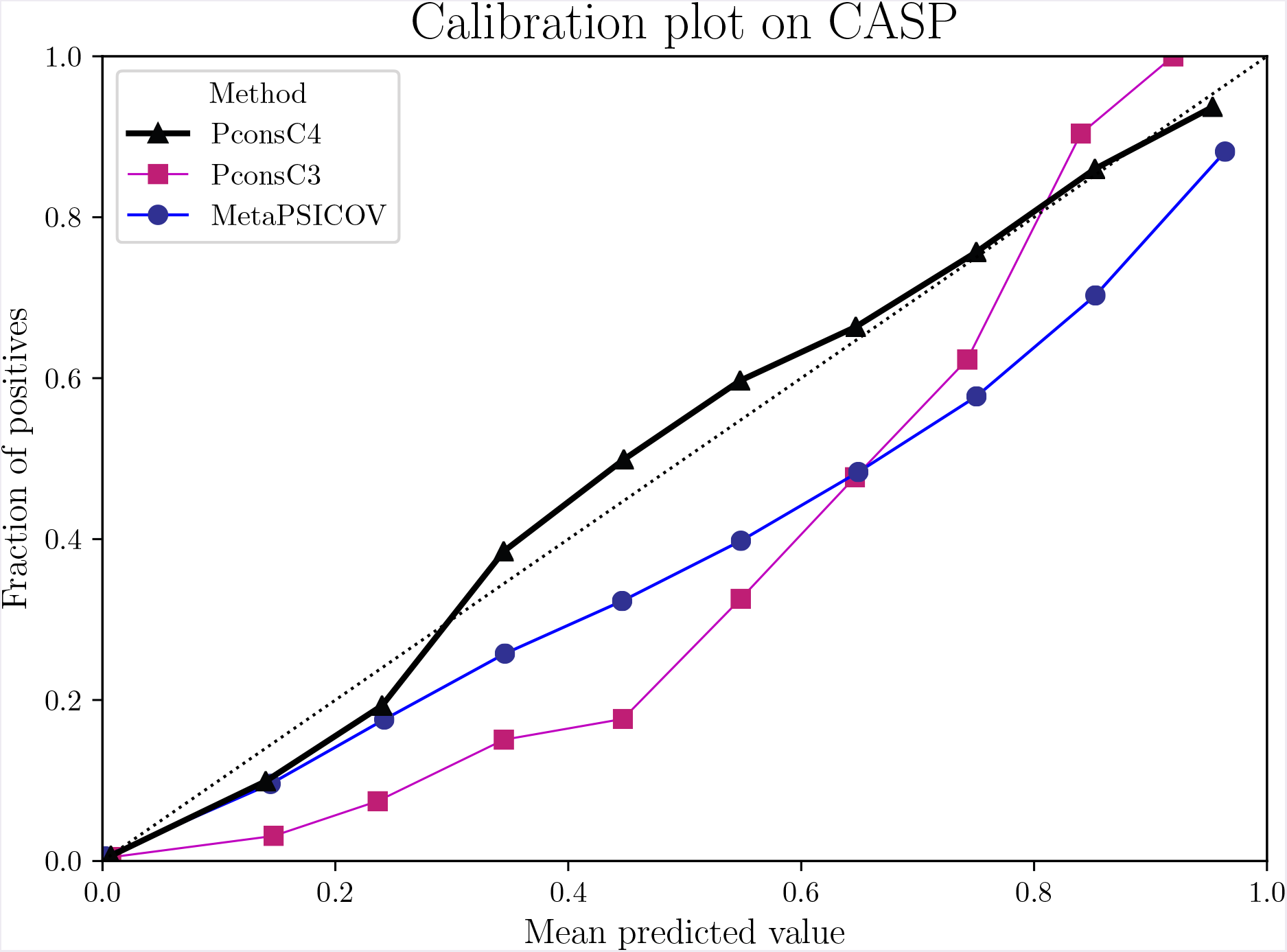
Calibration curves on CASP12

### 1.2 Precision as a function of the alignment depth

The Figure S5 shows performance vs. number of effective sequences. Here we use the same definition as in (Baldassi *et al.*, 2014a): the number of sequences that are significantly different from each other. While statistical methods, like DCA, benefit from a constant improvement for larger number of effective sequences, PconsC4 starts to plateau at around 10^2^ sequences. This suggests that it is capable of learning to ignore the noise in DCA methods, even when the signal is weak.

**Figure S5.**
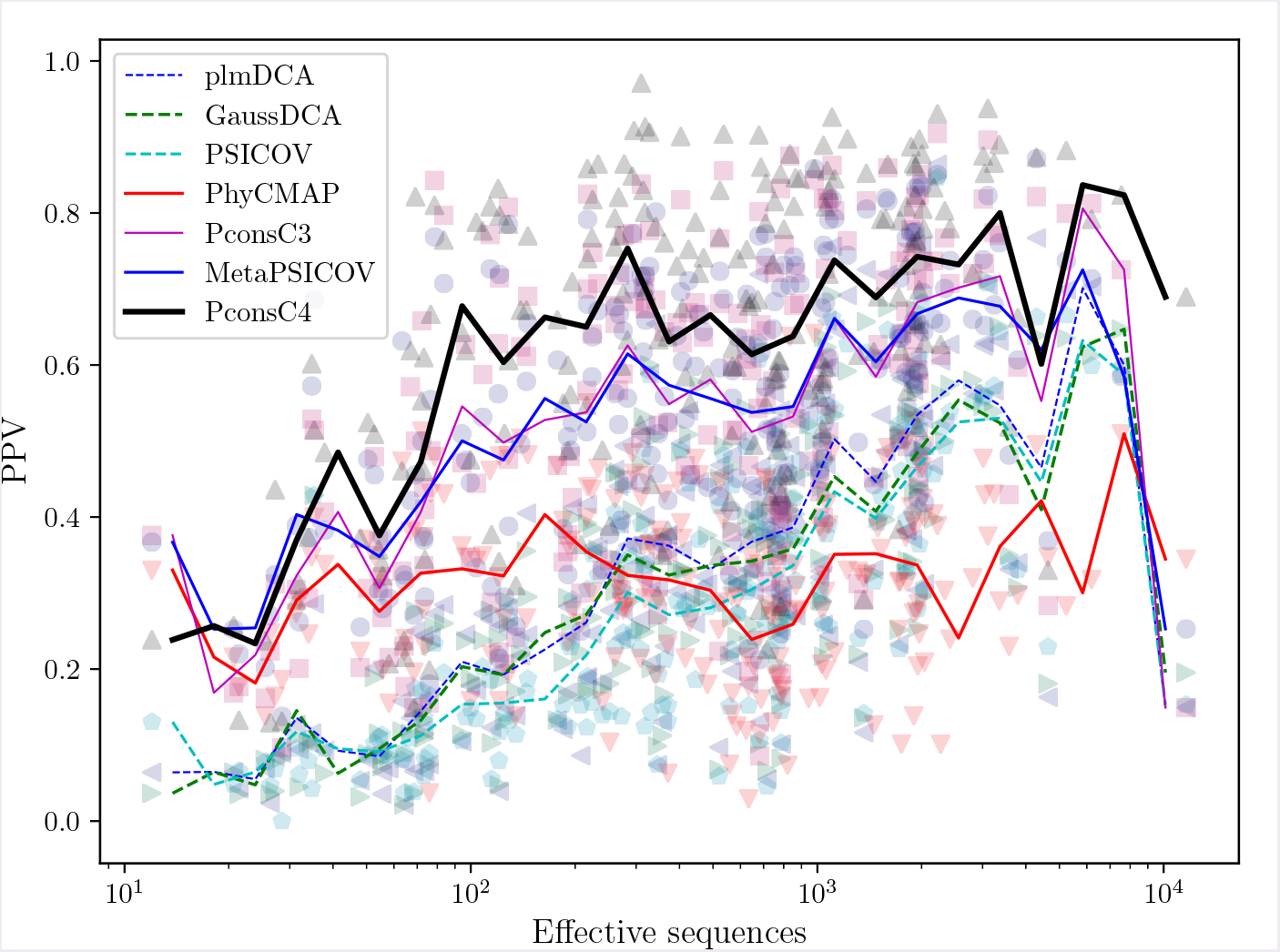
PPV vs. number of effective sequences on the benchmark set. The top *L* contacts are considered.

## 2 GaussDCA optimization

In the reimplementation of GaussDCA we made two differences with respect to the reference implementation that improve the speed, while keeping the results exactly the same.

### 2.1 GaussDCA and compressed alignments

Much of the CPU time is spent computing sequence weights, which means we need to compute all pairwise Hamming distances between sequences in the multiple sequence alignment. In a naive implementation, we store the alignment as a 2D array of int8. The compiler can then use SIMD instructions to combine several numbers to fit them in the word size of the CPU.

The Julia implementation of Gaussdca, (Baldassi *et al.*, 2014b), uses a manually compressed alignment: since we have only 21 states, we only need 5 bits per symbol, so we can manually pack up to 12 amino acids in a single 64 bits int, that fits our CPU optimally. To compute the distance, we need to XOR the sequences, and count the number of 5-bit regions where it differs. This gives us a 50% increase in packing efficiency with respect to the naive 8-bit storage.

In our re-implementation we found that the naive version is actually faster, probably because it is easier for the compiler (GCC 7) to optimize. Further improvements can be obtained activating auto-vectorization (GCC flag `–ftree–vectorize`), suggesting that the code emitted by Pythran is more suitable for vectorization.

### 2.2 Fast estimation of the expected similarity threshold,*θ*,in GaussDCA

Two sequences are considered similar if their Hamming distance is below a given threshold. (Baldassi *et al.*, 2014a) introduced an automatic estimation as the average fraction of differences across the alignment, and implemented it by directly counting them, which is 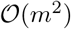 on the number of sequences in the alignment.

The same result can be obtained in linear time by counting the number of occurrences of each symbol in each column (called the bincount function), and computing the 2-combination 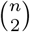. For an alignment of *C* columns and an alphabet of size *Q*:

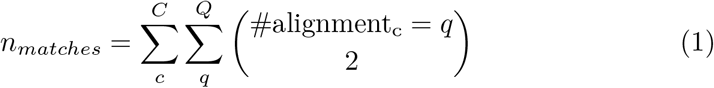

## 3 Training

### 3.1 Description of inputs

#### 3.1.1 One-dimensional inputs

From the columns of the multiple sequence alignment we can compute the probability of finding each amino acid or gap, *p¿.* For amino acids, we compute the average frequency 〈*pi*〉 as the observed frequency of the amino acid on the Uniref50 dataset. The expected probability of gap 〈*p*_ 〉 is estimated from each alignment. We consider a total of 23 amino acid states, the usual 20, plus the gap state, plus B (asparagine or aspartic acid) and X (unknown).

For each column we can compute self-information as the vector of entries:

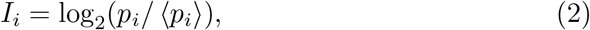

and the partial entropies are defined as:

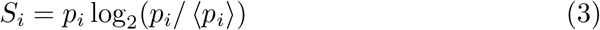

The sequence is given as one hot encoding.

#### 3.1.2 Two-dimensional inputs

The two-dimensional inputs are mutual information (MI), normalized mutual information (NMI), cross entropy (H), and GaussDCA.

Mutual information is defined as:

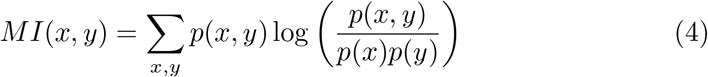

We also include two more inputs that are adjusted versions of MI. The first is normalized mutual information, where we think of *MI* as an analogue of a covariance, and *NMI* is the Pearson correlation coefficient. The entropies of the columns play the role of the variances:

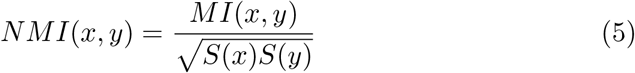

And lastly, cross entropy is calculated as an additive normalization of mutual information:

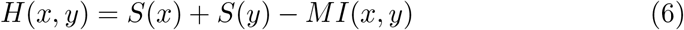

The Average Product Correction (APC) (Dunn *et al.*, 2008) is applied to all two-dimensional inputs except for cross entropy.

### 3.2 Training and testing sets

PconsC4 is trained on a set of2891 proteins culled from PDB using PISCES (Wang and Dunbrack, Jr., 2003) with a maximum sequence identity 20%, minimum resolution 2.0 Å, maximum R-factor 0.3. Furthermore, chains from the same ECOD (Cheng *et al.*, 2014) H-group as any protein in the benchmark dataset or dating from after 2016-05-01 was removed to avoid potential overlap with the test datasets. Out of these 2891 proteins, 100 randomly selected proteins were used as a validation set for optimization, see Table S3 and S4. For benchmarking, two datasets are used, the same 180 proteins, Table S5 as in (Michel *et al.*, 2017) and the 44 proteins from CASP12 with available structures, Table S6

## 4 Pipeline

We include a schematic of the complete pipeline, see Figure S6.

In the upper left corner are the 1D inputs being fed to the pre-trained model taken from ProQ4 (Menéndez Hurtado *et al.*, 2018). It is pre-trained to predict secondary structure and surface accessibility for each residue (golden outputs in the middle left).

The 128 output channels of the second to last layer are used to extract relevant 1D features. These are then transformed into 2D features by the outer product and combined with the remaining 2D features (GaussDCA, Mutual Information, Normalized Mutual Information, and Cross Entropy). The concatenation is then passed on to the U-net block (lower rectangle).

U-net was developed for image segmentation and combines convolutional layers, max pooling (downwards arrows) to increase the effective receptive field, upsampling (upwards arrows) to recover the original size, and skip connections (horizontal arrows) to help convergence.

The final predictions are shown on the upper right: contact maps at the three distance thresholds and distance as S-score. The branch predicting S-score is solving a regression problem with a Mean Squared Error loss, whereas the other three output branches are classifying the probability of a contact for the thresholds of 6, 8, or 10 Å, and use a cross entropy loss.

All intermediate convolutional layers are followed by a ELU (Clevert *et al.*, non linearity, Batch Normalization (Ioffe and Szegedy, 2015), and a Dropout layer (Srivastava *et al.*, 2014) with a probability of 0.1. Weights are initialized using the He distribution as described in the ResNet paper (He *et al.*,2016). A weight decay of 10^−12^ on the *L*^2^-norm is applied for regularization. All output layers have an appropriate function to ensure the output is in the valid range, sigmoid in our case.

The model is trained for 100 epochs (full passes over the training data) using the Adam optimizer at an initial learning rate of 0.001. The learning rate is decreased by a factor of 0.5 if the S-score training loss did not decrease over the last five epochs. The training dataset is shuffled after each epoch. The final model was selected by taking the epoch with minimum S-score loss on the validation dataset (epoch 29).

**Figure S6.**
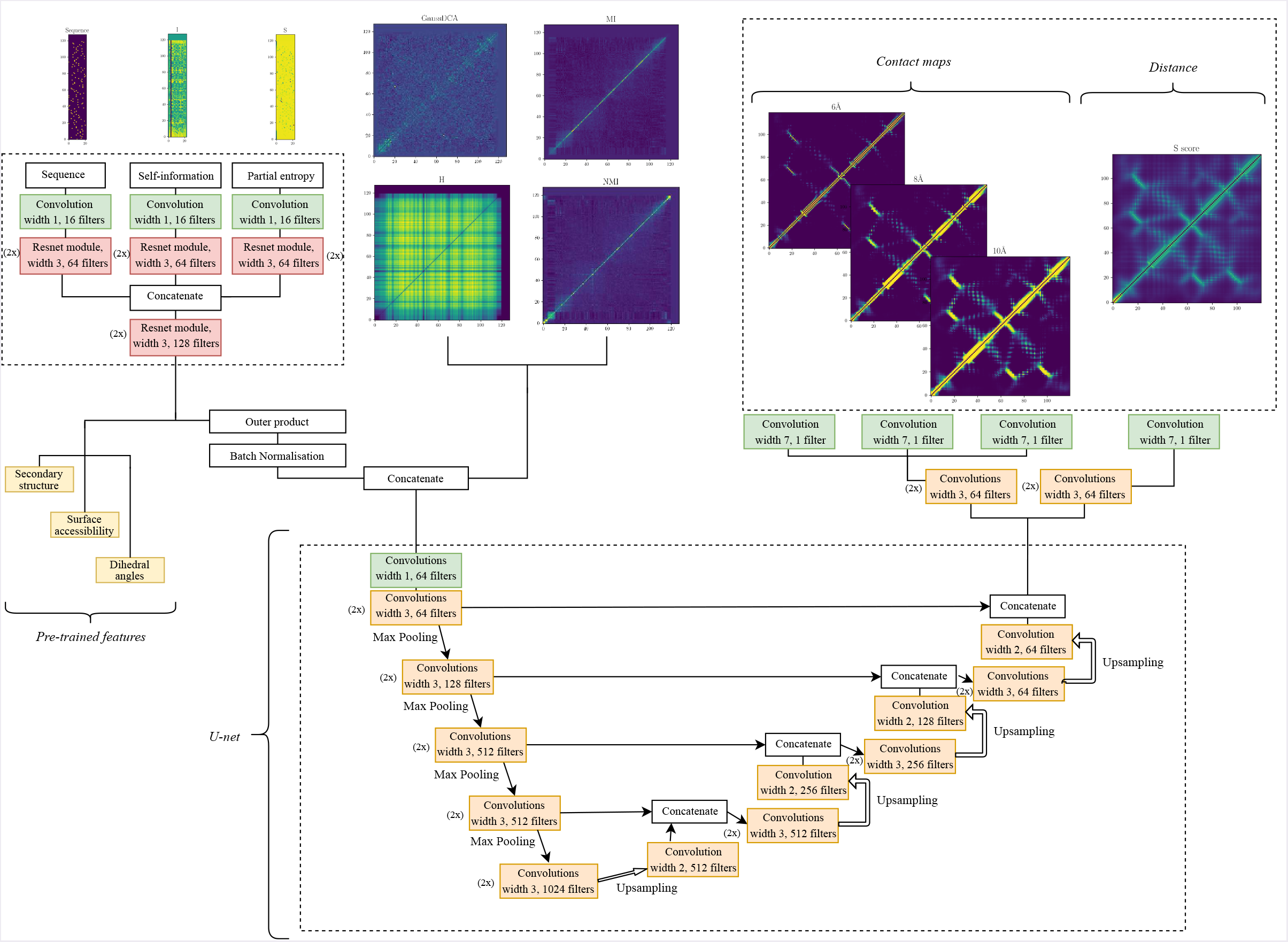
The example is the protein 2AV5 chain D from the benchmark set.

Figure S6: The example is the protein 2AV5 chain D from the benchmark set.

## 5 Instructions

### 5.1 Installation

On a machine with Python 3.5 or higher and pip, PconsC3 can be installed using the following commands:

pip install numpy Cython pythran wget https://github.com/ElofssonLab/PconsC4/releases/download/0.2/\pconsc4–0.2.tar.gz

`pip install pconsc4-0.2.tar.gz`

Once additional space is granted in PyPI (Python Package Index), a simple pip install pconsc4 will be sufficient.

A deep learning backend compatible with Keras will also be needed. We recommend Tensorflow:

`pip3 install -U tensorflow`

### 5.2 Example usage

`import pconsc4`

`# Load the deep learning model model = pconsc4.get_pconsc4()`

`# It is possible to re-use on different alignments pred_1 = pconsc4.predict(model, ′path/to/alignment1′) pred_2 = pconsc4.predict(model, ′path/to/alignment2′)`

`# Show pred_1 on the screen: import matplotlib.pyplot as plt plt.imshow(pred_1[′contacts′][′cmap′]) plt.show()`

`# Save pred_2 in CASP format: from pconsc4.utils import format_contacts_casp print(format_contacts_casp(pred_2[′contacts′][′cmap′], seq_2, min_sep=5))`

### 5.3 Supported formats

The program accepts alignments in. `fasta, .a3m, or .aln`, without line wrapping.

## 6 Training and test sets

PconsC4 is trained on a set of 2891 proteins culled from PDB using PISCES (Wang and Dunbrack, Jr., 2003) on 2017-09-14 with the following criteria: maximum sequence identity 20%, minimum resolution 2.0 Å, maximum R-factor 0.3. Furthermore, chains from the same ECOD (Cheng *et al.*, 2014) H-group as any protein in the benchmark dataset or dating from after 2016-05-01 was removed to avoid potential overlap with the test datasets. Out of these 2891 proteins, 100 randomly selected proteins were used as a validation set for optimization.

**Table S3.**
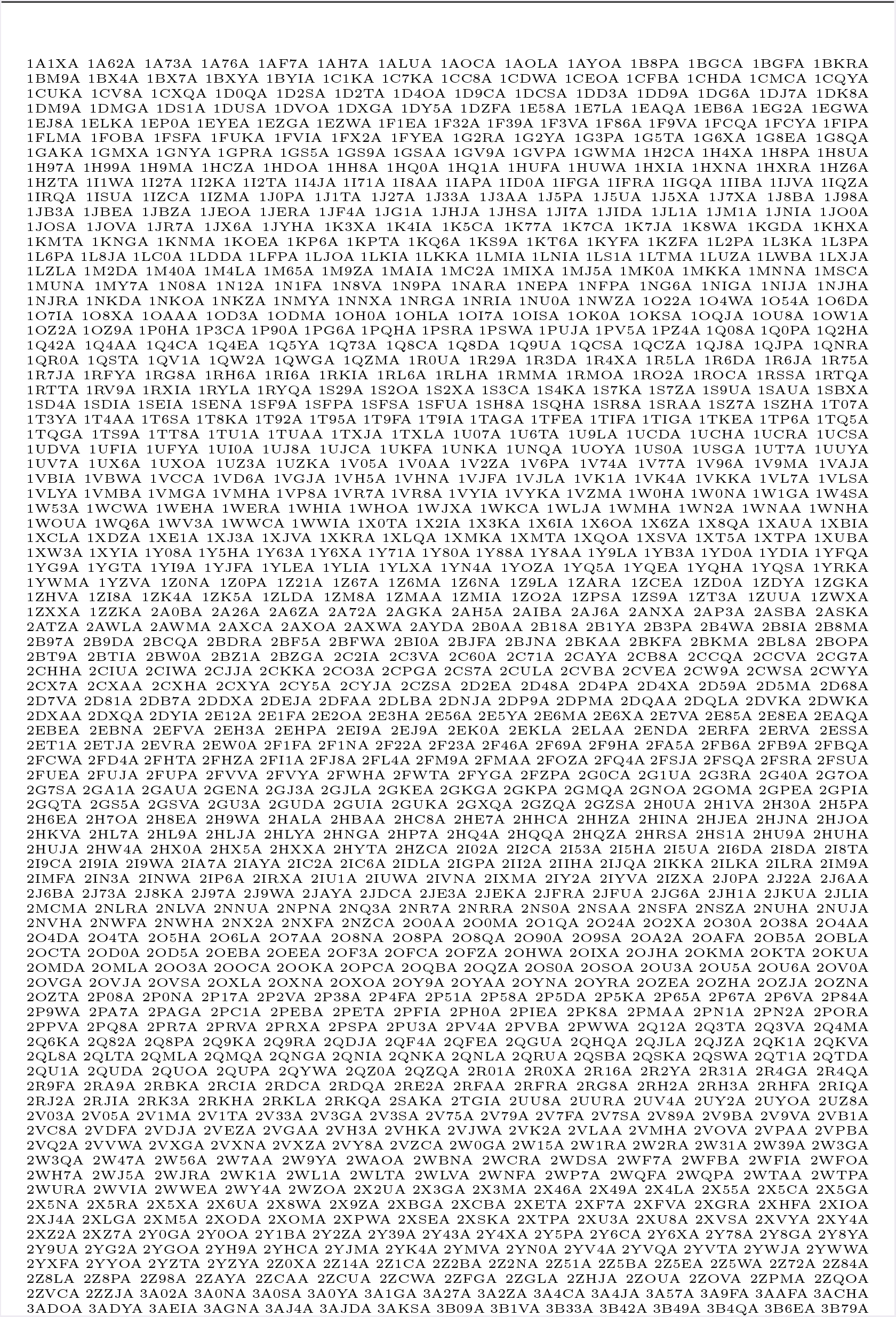
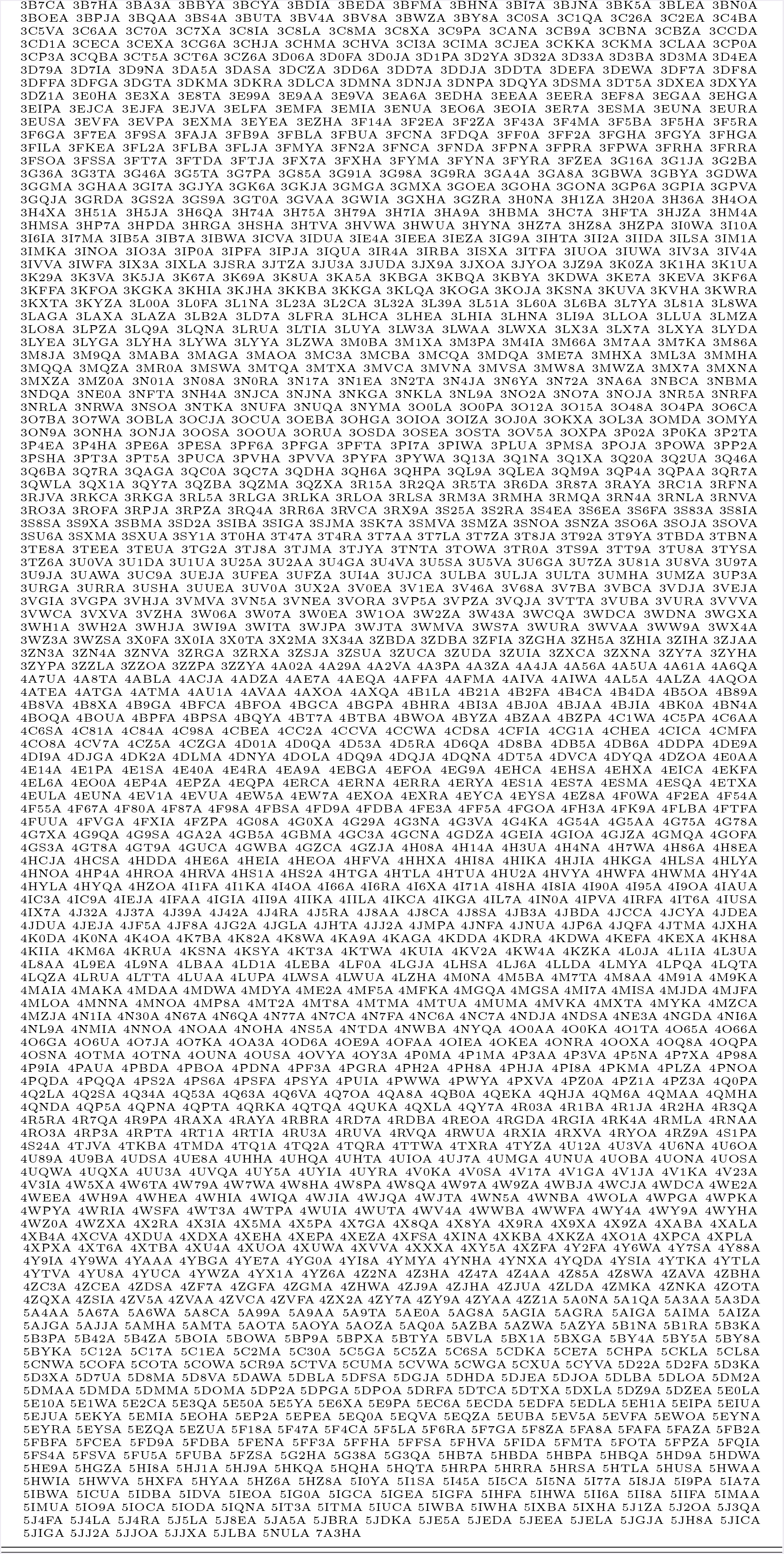
PDB code and chain identification for proteins in the training set

**Table S4.**
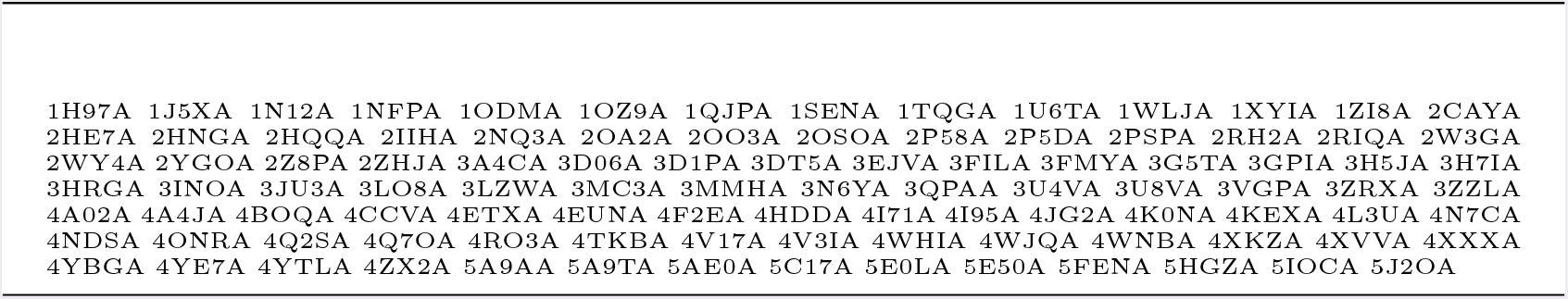
PDB code and chain identification for proteins in the validation set

**Table S5.**
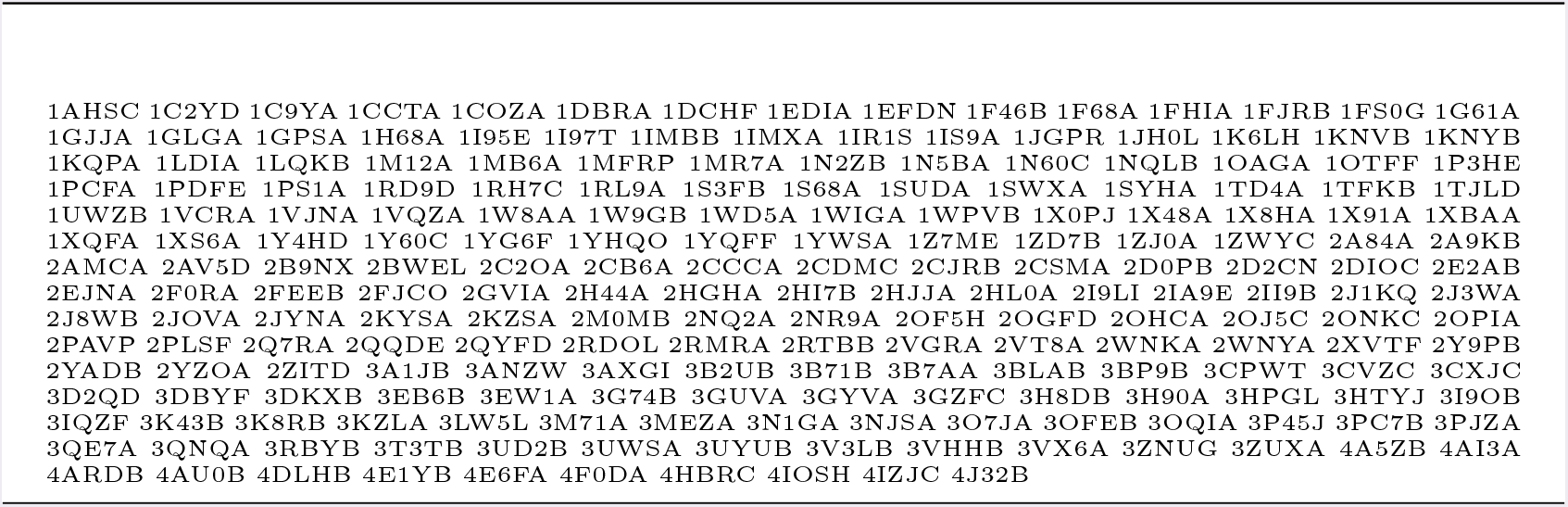
PDB code and chain identification for proteins in the benchmark set

**Table S6.**
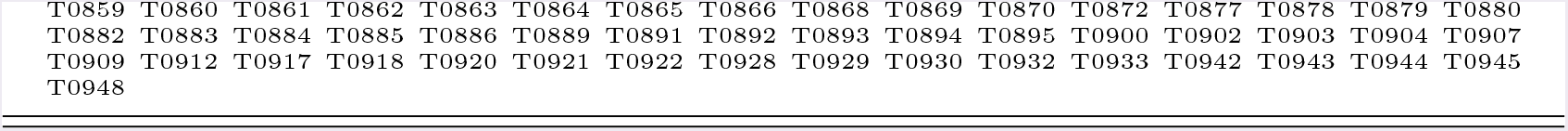
PDB code and chain identification for CASP12 targets

